# The SSVEP tool as a marker of subjective visibility

**DOI:** 10.1101/588236

**Authors:** Adélaïde de Heering, Arnaud Beauny, Laurène Vuillaume, Leila Salvesen, Axel Cleeremans

**Affiliations:** CO3, Centre for Research in Cognition & Neurosciences, Université libre de Bruxelles, Avenue F. Roosevelt, 50 CP 191, 1050 Bruxelles, Belgium

**Author notes:** Correspondence to: Adélaïde de Heering, Centre for Research in Cognition & Neurosciences, Université Libre de Bruxelles, Avenue F. Roosevelt, 50 CP 191, B-1050 Belgium.

**Keywords:** consciousness, subliminal, faces, steady-states, no-report

## Abstract

This study is part of a larger attempt to explore how the brain produces conscious experience. Our main objective here was to take advantage of a neural signature conveyed by the steady-state visual evoked potentials (SSVEP) technique (1) to explore the extent to which complex visual images can be processed in the absence of consciousness and (2) to determine whether this tool can be used to shed light on participants’ phenomenal experience of these images. To this end, we embedded faces within sequences of non-face stimuli and manipulated their contrast to create a subliminal condition. Our results were threefold. First, we show that a significant brain activation can be delineated with the SSVEP tool even when participants report being unable to see the stimuli. In this subliminal condition, the brain response was confined to the back of the scalp. Second, we observe that the face signal increases in magnitude and propagates bilaterally along a posterior-to-anterior axis as image contrast increases. Third, we suggest that SSVEP could be used as a novel instance of a no-report paradigm because it requires no overt behavioural response, and because at some contrast levels and electrodes, its outputs (signal magnitude and scalp topography) predict people’s self-reported phenomenal experience.

## INTRODUCTION

Identifying the neural correlates of conscious experience remains a significant challenge in the field of cognitive neurosciences. Since the seminal work of Crick and Koch [1], it has become clear that the very notion of a “neural correlate of consciousness” is problematic. It is unclear, for instance, whether we are looking for a single area, for a network of interacting areas, or for a specific kind of process underlying conscious experience [2]. Likewise, it is still unclear how to best distinguish the “true” neural correlates of consciousness from its prerequisites (NCC-pr) and from its consequences (NCC-co) - in other words: how do we neutralize what is specific to conscious experience from other factors such as working memory or selective attention that are known to contaminate the identification of the specific mechanisms that underpin conscious experience [3-4-5]? Furthermore, multiple confounds have been identified that also compromise this endeavour. Lau & Passingham [6], for instance, showed that performance, which is typically better when people report seeing a stimulus than when they do not, elicits its own neural correlates. Similarly, the typical requirement for participants to report on their conscious state also generates report-driven activity in the brain that may not be observed when people are not asked to produce them [7; see also 5, 8-9]. Lately, these concerns have motivated the development of so-called *no-report paradigms*, which offer the promise of building indirect measures of subjective experience, hence making it possible to avoid verbal or motor reports. Technically, such paradigms also offer the opportunity to get closer to real-life situations, during which individuals seldom reflect, or report, on their thoughts or experiences of the world. Thus far, amongst possible indirect measures are simple physiological measures, such as eye movements or pupil dilation/constriction, the modulations of which have been used to objectively and continuously map perceptual alternations from binocular rivalry stimuli, for example [10; for a review see 11]. Invasive local field potentials (LFP; [12]) and general flash suppression (GFS; [13-14]) have also been used to shed light on which brain regions participate to subjects’ phenomenology. Most of these measures nevertheless lack sensitivity (e.g., all-or-none responses) and flexibility (e.g., the requirement of keeping participants’ head fixed).

In this context, the major goal of this study was to explore whether a non-invasive technique derived from electroencephalography (EEG) – the steady-state visual evoked potentials (SSVEP) method [17] – could be used to access participants’ phenomenology. To test so, we relied on a sensitive paradigm originally designed to capture participants’ brain responses when they form a category from very different visual stimuli (such as faces) and exclude the exemplars that do not belong to this category [15-16]. Technically, it only requires that participants view sequences of images flickering at a given frequency (e.g., 6 Hz = 6 images/second = image frequency) while their EEG is recorded (for evidence on pre-motor and pre-verbal infants with a similar paradigm, see [18]). In the current study, images were organized so that a face would re-appear every 5^th^ item in a 6 Hz sequence, thus, at the slower frequency of 1.2 Hz (6 Hz/5 = face frequency). Crucially, visibility, indexed through overall contrast, was equally degraded for all the images composing a given sequence, and also steadily increased over the course of EEG recordings. As such, we expected participants not to anticipate faces to re-appear every 5^th^ item in the low-contrast conditions given that they were kept unaware of any kind of periodicity contained in the SSVEP sequences. In addition, the experimental design allowed us to address other important questions such as: Is it possible to identify the neural correlates associated with exposure to stimuli degraded to the point that participants report not having seen them? Does this capability extend to the complex process of forming a visual category from very different items? Where does such processing happen in the brain, and how does this activity propagate when the stimulus becomes visible to participants?

Given previous findings [19], we hypothesized that the face categorization paradigm we used here would be sensitive to address such questions, even at subliminal stages - not only because of the documented power of the technique, but also because faces are stimuli that are both very salient and familiar to people. This should undoubtedly constitute an advantage in any attempt to uncover the brain regions activated during subliminal processing [20]. We therefore expected to find significant brain activation at the frequency at which faces were introduced within the sequences, namely at 1.2 Hz (and harmonics), and even at a contrast level defined as subliminal for the group of participants. We also assumed this subliminal activation to be located at either occipito-temporal electrodes lying relatively near to these cortical regions known as being particularly sensitive to faces [15-16], but thenceforth to a lesser extent than what is typically observed at high contrast levels, or at more posterior electrodes (e.g., the occipital lobe; [21]). Finally, we predicted, in line with previous studies (see [22] for a review), that the SSVEP tool would be sensitive enough to track down, over the course of only a few trials, the involvement of different electrodes, on a posterior-to-anterior axis, as the overall contrast of face images increases.

Relating this SSVEP paradigm to participants’ phenomenal experience (“what it feels like to see a face”; [23]) is a task harder to achieve because it does not simply require showing that low- and high-contrasted stimuli, and their underlying brain regions, are associated with low and high visibility ratings. This indeed would be totally uninformative to document on what their conscious experience is like, given that high-contrast stimuli are typically rated as more visible than low-contrast stimuli [24-25]. We therefore proceeded with what is typically performed in binocular rivalry paradigms [26-29] for which emphasis is placed on the subject’s differential phenomenology when stimulation is kept constant and investigated whether we could find any meaningful relationship between the objective and the subjective measurements collected during the EEG session. To this end, we inspected each contrast level and visibility scale separately and expected the correlation/regression equations we computed for this purpose to be particularly informative about this relationship.

## METHODS

### PARTICIPANTS

The Research Ethics Board of the department of Psychology of the Université libre de Bruxelles (Belgium) approved all experimental protocols (Reference - B406201734083). All methods were carried out in accordance with relevant guidelines and regulations. We tested 15 French-speaking adult participants (5 males; all right-handed and with normal or corrected-to-normal vision; mean age: 22 years, SD = 3; one participant excluded because of high artefacts on more than 2 EEG sequences/block). All gave written informed consent and received financial compensation in exchange for their participation.

### STIMULI

We used a previously described set of 200 × 200-pixel coloured images [18; see also 15-16]. This set contained 48 face images and 248 object images (e.g., animals, houses, fruits/vegetables, flowers/plants, man-made objects). Face images varied in terms of colour composition, age, sex, viewpoint, lighting conditions, and background (for an example of the stimulus set used in the current study, see [18]). After equalizing the overall contrast and luminance of all images, we used Adobe Photoshop CS6 to create 11 image sets of varying visibility by parametrically manipulating the contrast of each image from 0% (no information) to 5% (some information), in 0.5% increments. All 11 sets were used for SESSION 1 (behaviour). We selected 5 out of these 11 sets (0%, 0.5%, 1%, 1.5%, 2% and 100% contrast) for SESSION 2 (EEG) based on behavioural measurements. Images used at SESSION 2 were slightly larger (6.7 × 6.7 degrees of visual angle) than those used at SESSION 1 (5.7 × 5.7 degrees of visual angle).

### PROCEDURE

The experiment included an initial behavioural test (SESSION 1) followed by an EEG component (SESSION 2). These sessions were scheduled to take place either on the same day or on two consecutive days and were not counterbalanced across participants (i.e., behavioural testing always preceded EEG recording). On both testing days, participants sat comfortably at a distance of 50 cm from a Lenovo laptop computer (refresh rate = 60 Hz). Stimuli were always presented in the centre of the screen and displayed by means of Matlab (The Mathworks).

<u>SESSION 1</u> (behaviour) was designed to yield both an objective and subjective index of the categorization threshold for faces. To this end, we presented each of the 48 faces twice, along with an equal number of object images, for a total of 192 randomized trials. For each briefly presented and unmasked image (83.5 ms duration), participants performed two tasks. The first task was a two-alternative forced-choice task (2AFC) for which participants had to categorize the image as either a face (FA) or a non-face (NF). This task was controlled through a one up/one down staircase procedure (initial contrast level = 3.5% contrast), which aimed at precisely identifying which contrast level the group needed to categorize a face as a face and differentiating faces from non-faces at exactly chance (50%) level. This threshold was also used to define the subliminal threshold used at SESSION 2. For the second task, participants were asked to give a subjective rating of the visibility of each image on a four-point version of the “Perceptual Awareness Scale” (PAS) [30-31]. More specifically, they had to choose between (0) no impression of the stimulus (no experience), (1) a feeling that something has been shown, with a content that cannot be specified any further (brief glimpse), (2) an ambiguous experience of the stimulus, with some aspects being experienced more vividly than others and a feeling of almost being certain of it (almost clear experience), or (3) a non-ambiguous experience of the stimulus with no doubt about its own answer (clear experience). Responses at both tasks were not speeded, i.e., participants could take as long as they needed to respond. Statistical tests included measures of sensitivity (d primes), *t*-tests applied with Bonferroni correction when necessary and repeated measures ANOVAs.

<u>SESSION 2</u> was dedicated to EEG recordings. Here we presented the same participants with the same images as at SESSION 1 even though they were arranged this time within streams of stimuli to meet the requirements of the SSVEP stimulation. More specifically, participants viewed series of images flickering against a grey background (image presentation frequency = 6 Hz, i.e., 6 images/second). Each image appeared on screen for just 83.33 ms, and was followed by a blank interval of 83.33 ms, for an image cycle duration of 166.66 ms. Contrary to Jacques and collaborators [15] which study inspired this one, we used a square-wave contrast modulation. Contrast varied therefore from 0% to the maximum in one cycle, the goal being to precisely control the amount of contrast participants were exposed to during a given sequence, and to guarantee that the presentation duration of each image precisely corresponded to that used during the behavioural session (i.e., 83.33 ms). Within each stream, a face image was embedded at every 5th item, which ensured the face-presentation frequency to be at exactly 6 Hz/5 and consequently, that a face-selective neural activity will be evoked at a pre-specified point in the EEG frequency spectrum, namely at 1.2 Hz. The rest of the sequence was composed of non-face stimuli. Every SSVEP sequence started with a “ready?” message presented on a uniform grey background that invited participants to press the spacebar to initiate the sequence. Once launched, a central black fixation-cross appeared for between 1 to 5 seconds, after which the stimulation sequence (43.33 s) began. A SSVEP sequence consisted of a 1.67-second fade-in period, a 40-second period of interest (48 faces, 192 non-faces) and a 1.67-second fade-out period. During the fade-in and fade-out, the maximal contrast of images progressively ramped up and down, respectively, so as to minimize blinks and artefacts elicited by the sudden appearance/disappearance of flickering stimuli. The previously mentioned fixation-cross remained superimposed on the images throughout the sequences. Participants’ task was to fixate it and to press the spacebar whenever it turned red. Importantly, this change would happen non periodically. We used this task, orthogonal to the image content, to keep participants active during the task. Statistical analyses involved the calculation of the correct detection of this change of colour of the cross relative to target onset and was only considered if its detection occurred between 250 and 1500 ms following target onset.

To examine how the response to the SSVEP stimulation varied as a function of stimulus contrast and phenomenal experience, all participants completed 6 blocks presented in a fixed order, in which image contrast increased progressively. That is, participants transitioned from seeing images at 0% contrast (i.e., dark grey squares flickering on a light grey background – Block 1) to images at 0.5% contrast (Block 2) immediately followed by images at 1% (Block 3) and 1.5% contrast (Block 4). The experiment ended with the presentation of images at 2% contrast (Block 5) and 100% contrast (Block 6). In this way, information gleaned by participants during the high-contrast conditions (e.g., that the faces appear at regular intervals in the sequence) could not influence the response captured during the lowest contrast conditions and everyone would endure the undetermined contamination effect between conditions to the same extent. Each SSVEP block contained 10 sequences.

Additionally, we assessed participants’ subjective impression about the images of each SSVEP sequence by collecting their ratings on two visibility scales in addition to what had been performed at SESSION 1 with the PAS scale. We did so mainly because we evaluated that phenomenal experience of sequences of images could be very different from the one of single images. We therefore first asked participants to judge whether among the grey square images they saw during the preceding sequence, they had (0) no impression of the stimuli (“you could not categorize any” – no experience), (1) brief glimpses of the stimuli (“you could categorize some of them” – brief glimpse), (2) an almost clear experience of the stimuli (“you could categorize almost all of them” – almost clear experience) or (3) a clear experience of the stimuli (“you could categorize all of them” – clear experience). Then we asked them to estimate how many of the 240 images they just saw they felt they could categorize (quantitative scale). In both cases, these ratings were used as a proxy of face visibility that we did not assess *per se* because we did not want to generate any expectation regarding this specific visual category, which could have contaminated the SSVEP observations. The analyses of these ratings involved repeated-measures ANOVA corrected for multiple comparisons (Bonferroni correction).

#### EEG recordings and analyses

We recorded scalp EEG using a 64-channel Biosemi ActiveTwo system (Biosemi, Amsterdam, Netherlands). The EEG analog signal was digitized at a 1024 Hz sampling rate. During recording setup, electrode offset was reduced to between ±50 μV for each individual electrode by softly abrading the underlying scalp with a blunt plastic needle and insulating the electrode tip with saline gel. Eye movements were monitored using 4 electrodes placed at the outer canthi of the eyes and above and below the right orbit.

EEG analyses were carried out using Letswave 6 (https://github.com/NOCIONS/letswave6) and custom scripts running on Matlab (The Mathworks). Importantly for the purpose of the current study, 5 regions-of-interest (ROIs) were delineated according to previous literature on the topic [see for example, 15; 21] and included 2 to 3 channels: ROI 1 (Iz-Oz), ROI 2 (O2-PO4-PO8), ROI 3 (O1-PO3-PO7), ROI 4 (P6-P8-P10) and ROI 5 (P5-P7-P9). Whereas noisy electrodes were linearly interpolated using 2 immediately surrounding clean channels (performed for 13 participants, with a maximum of 4 interpolated channels per participant), electrodes belonging to these ROIs were interpolated in reference to the other electrodes of that same ROI. No interpolation involved the Iz or Oz channel. Epochs were re-referenced to a common reference computed using the average of all 64-scalp channels (excluding ocular channels) and further segmented to contain an exact integer number of 1.2 Hz cycles beginning 2 seconds after the onset of the sequence (i.e., after the fade-in period, at the start of the proper sequence) until approximately 40 seconds after sequence onset (i.e., immediately before the fade-out period). For each participant, we created conditional averages for each of the 6 contrast levels (0%, 0.5%, 1%, 1.5%, 2% and 100%) and subjected the resulting averaged time series to Fast Fourier Transform (FFT), producing 6 frequency spectra (frequency resolution of 0.025 Hz) for each participant (one per condition). We also computed conditional grand-averages by averaging the FFT spectra across participants. In order to evaluate the magnitude of the effect across conditions and visualize it, two computations were performed on these files. The first computation involved the transformation of both individual and grand-averaged FFT spectra into signal-to-noise (SNR) ratios that were computed as the *ratio* between the amplitude at each frequency and the average of the 20 surrounding bins (10 on each side skipping the immediately adjacent frequency bins; for similar analyses see [15-16; 18; 32]). The second computation involved the transformation of grand-averaged FFT spectra into z-scores in a similar fashion, except that z-scores were computed as the *difference* between amplitude at the frequency of interest and the mean amplitude of the 20 surrounding bins divided by the standard deviation of the 20 surrounding bins. As in Jacques and collaborators [15], the z-score threshold was set at 3.09 (p < .001, 1-tailed) and was used to determine whether the signal at a given bin statistically differed from noise as well as the number of significant, consecutive and minimally involved harmonics across conditions that should be taken into account in the analyses. Beside these analyses, statistics also involved correlation/regression analyses, which main goal were to help deciding on whether the SSVEP outputs could serve as objective measures of subjective experience.

## RESULTS

Testing involved two sessions. The initial behavioural session (SESSION 1) essentially aimed at defining how contrast levels would be organized for the EEG experiment (SESSION 2). To this end, participants categorized briefly presented images (83.33 ms) as either a face (FA) or a non-face (NF) which contrast was adjusted using a one up/one down staircase procedure (range = 0-5%, increment = 0.5%). They also judged their visibility by means of a version of the “Perceptual Awareness Scale” (four-point scale PAS; [30]). The same participants’ scalp EEG was then recorded at SESSION 2 while they viewed these images but this time displayed within sequences of stimuli. The EEG experiment was composed of 6 blocks and started with what had been defined at SESSION 1 as subliminal to participants. Critically, the contrast of images was fixed within a block, and increased systematically from one block to the next to avoid expectations being built over the course of the experiment. Each block contained 10 sequences in which images were always presented in continuous 6 Hz streams (i.e., 6 images per second) for 40 seconds, with a face image presented every 5^th^ item. After each sequence, participants were asked to report on the visibility of the images by means of two visibility scales.

### (1) Behavioural measurements

For SESSION 1, in which participants had to explicitly categorize single images as faces (FA) or non-faces (NF) (Figure 1A), the staircase converged over contrast levels ranging from 0.8% to 2.4%. The average convergence value for the group calculated over 192 trials was 1.45% (Figure 1B). Participants’ proportion of correct responses for all 11-contrast levels ranged between 0.48 (SE = .03) at 0% contrast and 0.88 (SE = .06) at 5% contrast (Figure 1C) and increased regularly as a function of image contrast (F(10,30) = 4.209, p < .001). Accordingly, the averaged d prime scores across contrast levels ranged between −0.03 and 0.87 and reached an averaged value of 0.37 (SE = .06; t(14) = 5.65, p < .001). The same (increasing) trend characterized participants’ PAS scores (F(10,30) = 12.04, p < .001). The lowest contrast level at which classification performance was statistically higher than chance was 2% contrast (one-sample t-test: t(14) = 2.704, p = .017). Amongst the conditions qualified by less contrast (< 2% contrast), we decided on the 0.5% contrast level as the subliminal one for SESSION 2, for conservative purposes. It was indeed the first level characterized by some signal (vs. 0% contrast) and, importantly, the PAS visibility ratings associated to this contrast level were also significantly different from the ones collected at 2% contrast (0.5% contrast: (“no impression of the stimulus”, range: 0-0.93, X = 0.28; SE = .08; 2% contrast: (“a brief glimpse experience”, range: 0.34-1.22, X = 0.80; SE = .09; t(14) = −7.254, p < .001; Figure 1D).

**Figure 1.**
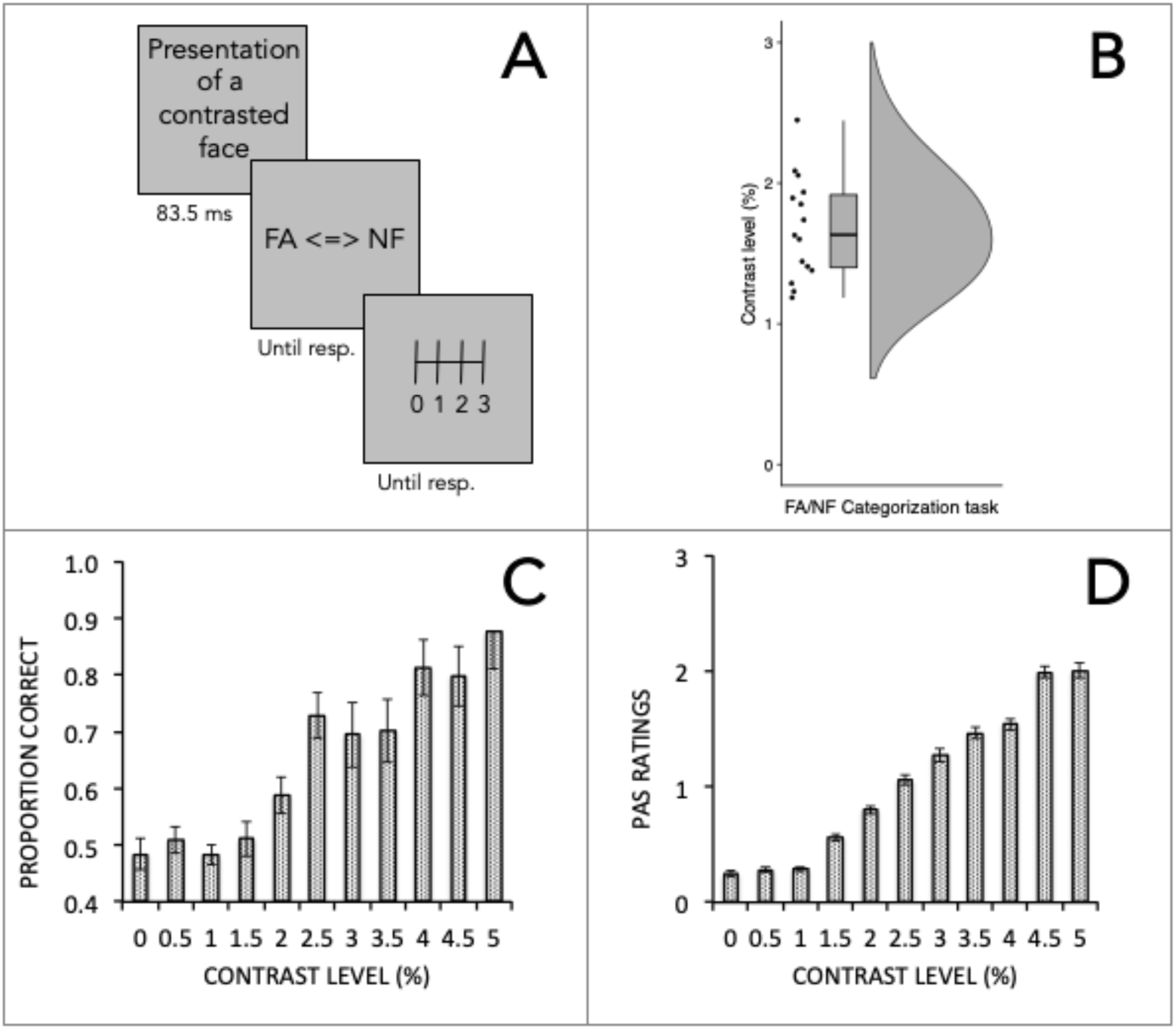
Behavioural definition of the subliminal threshold based on the data collected at SESSION 1. A. Participants (N=15) saw briefly presented images whose overall contrast level ranged between 0-5%. They categorized each as either a face (FA) or a non-face (NF) before rating image visibility on a 4-point PAS scale (0 to 3). B. The dots of the raincloud plot represent individual data. The thick horizontal line of the boxplot and the whiskers refer to participants’ averaged performance obtained at the staircase and its overall distribution. The staircase procedure identified 1.45% image contrast as the threshold for chance level (50% correct) at the FA/NF categorization performance. The far-right part of the raincloud plot should be interpreted as a density plot. C-D. There was a steady increase in the proportion of correct responses and in subjective visibility ratings as a function of image contrast. Bars and error bars refer to group performance and standard errors, respectively.

### (2) Preliminary exploration of the SSVEP data

For SESSION 2, participants were successively tested on 6 blocks varying in terms of the contrast of their images (0%, 0.5%, 1%, 1.5%, 2% and 100% contrast). These contrast levels were pre-selected from SESSION 1 and presented in a fixed order. Participants were all very accurate at the orthogonal task, namely at detecting the randomized colour-change of the fixation cross at each contrast level, even if their performance decreased at higher contrast levels given that the cross became somewhat less distinct from the background in these blocks (0%: X = 0.90, SE = .03; 0.5%: X = 0.85, SE = .05; 1%: X = 0.82, SE = .07; 1.5%: X = 0.67, SE = .07; 2%: X = 0.69, SE = .07; 100%: X = 0.69, SE = .08; F(4,52) = 6.569, p < .001). Importantly however, they allocated their attention to the fixation cross in a similar way at comparable contrast levels (e.g., 0% and 0.5% contrast), as indicated by the absence of significant differences between adjacent contrast levels (post-hoc Bonferroni tests: ps > .05). After each SSVEP sequence, participants’ subjective impressions of having seen the stimuli was also assessed by means of two visibility scales: a qualitative scale resembling the 4-point PAS scale and a quantitative scale through which participants had to report the number of images, up to 240, they felt they had been able to categorize within a given sequence. Participants carefully completed that task, distributing their responses across the full range of ratings. When all contrast levels were considered in the analysis, qualitative ratings significantly differed from each other (F(5,70) = 173.761, p < .001), as were quantitative ratings (F(5,70) = 96.594, p < .001). In both cases, the only significant difference recorded was between the adjacent 1% and 1.5% contrast levels as well as between the adjacent 2% and 100% contrast levels (Figure 2).

**Figure 2.**
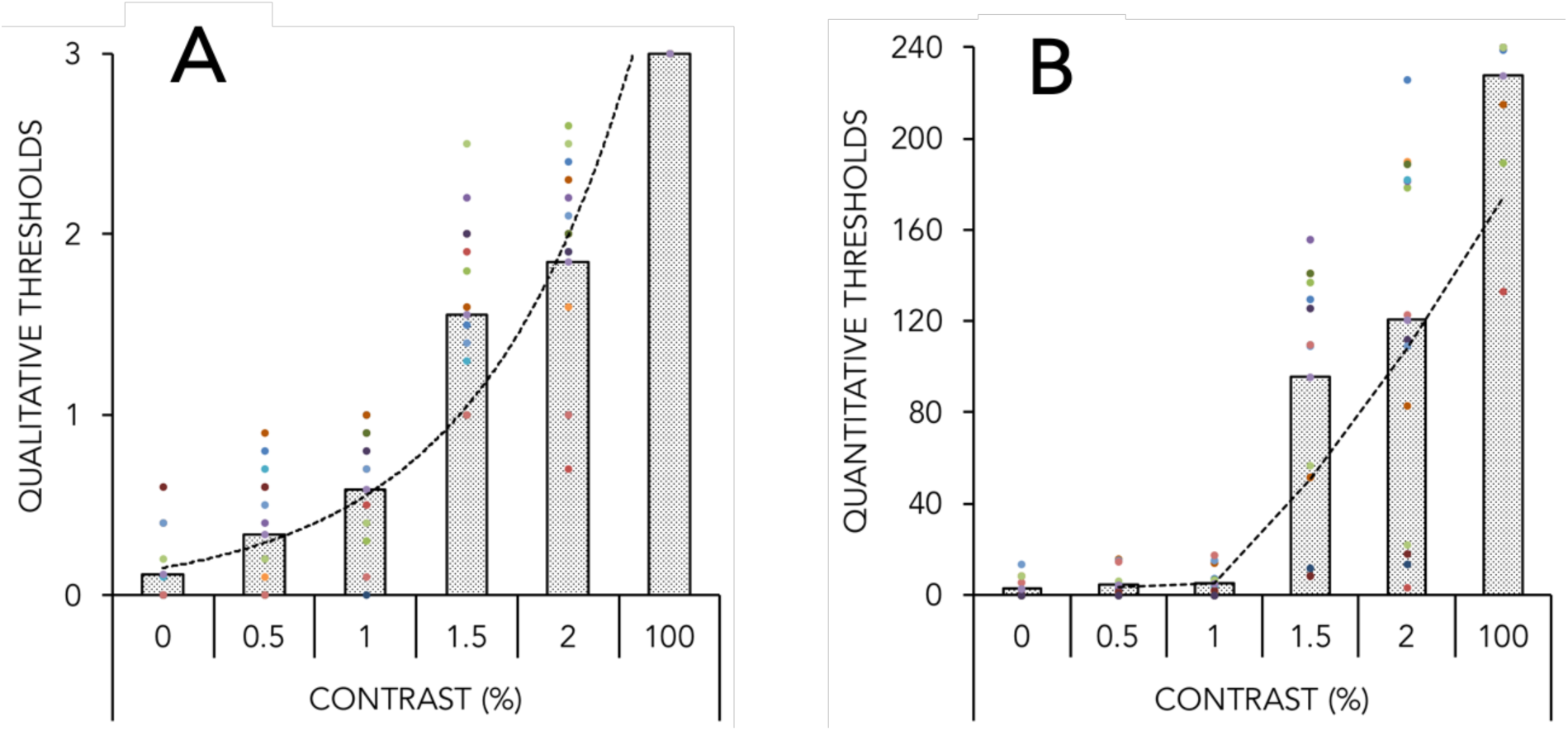
Subjective visibility ratings collected after each SSVEP sequence at SESSION 2 and further correlated with the SSVEP signal. There were two scales: one qualitative scale (A), which was inspired by the original PAS scale and one quantitative scale (B), which has been developed to mimic the characteristics of SSVEP stimulation (240 images presented within a sequence). Visibility ratings progressively increased with increasing contrast and followed distinct trajectories according to the scale, namely an exponential (A) and a stepwise distribution (B). Bars illustrate group performance. Dots illustrate averaged performance across sequences recorded at an individual level. Participants’ colour code is consistent between (A) and (B).

When images were presented at 100% contrast, that is, when they were completely visible to participants, we replicated the findings of previous studies [15-16] with a massive activation at the medial occipital lobe, centred on Oz, in response to the succession of the 6 images per second, which spread significantly over 4 consecutive harmonics (averaged brain SNR at 6 Hz and harmonics extracted at Oz at 100% contrast = 12.96 (SE = 1.39); Figure 3). Such responses typically indicate visual processes that are common to both faces and non-faces. The wave distribution and scalp topography extracted for images not contrasted at all (0% contrast) resembled the one extracted at 100% (i.e., activation centred on Oz and involving multiple harmonics). The magnitude of the peak at 0% contrast was however much smaller (averaged brain SNR at 6 Hz and harmonics extracted at Oz at 0% contrast = 8.16 (SE = .73)) than the one at 100% contrast because in the former case, participants only experienced a grey square flickering on a lighter grey background (paired t-test performed at Oz: t(14) = 3.646, p = .003). As in other studies [33], peak magnitude at 6 Hz and harmonics remained centred on Oz and monotonically decreased with decreasing discernibility across blocks (from 100% to 0% contrast: F(5,70) = 13.76, p < .001).

**Figure 3.**
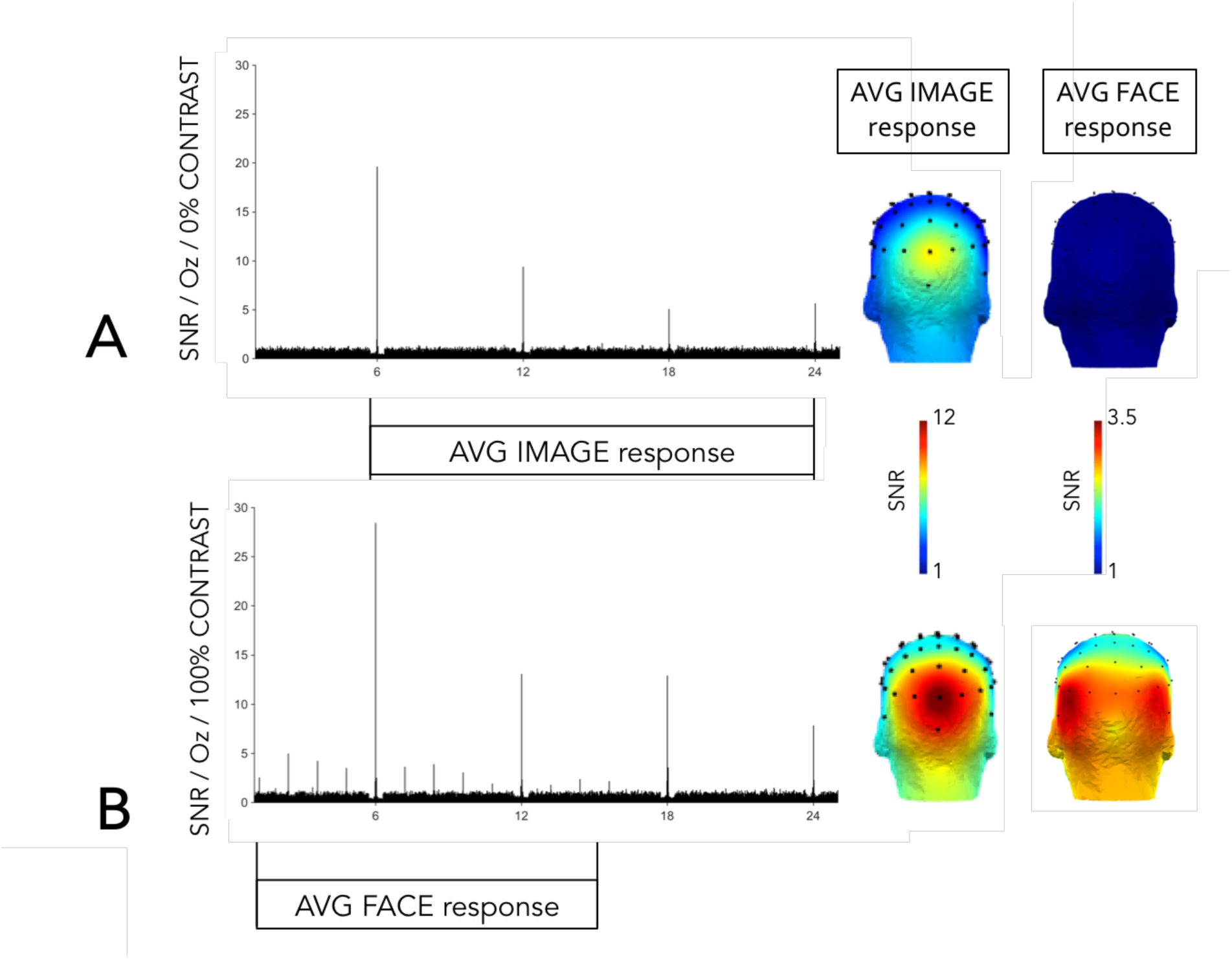
Signal-to-noise (SNR) spectra and averaged scalp topographies across significant harmonics generated at 0% (A) and 100% contrast (B) in response to face images. The SSVEP experiment consisted of images being repeated at the rate of 6 Hz (6 images/second) and of faces presented at the rate of 1.2 Hz (1 face image out of 5 items). Both the 0% and 100% contrast conditions lead to significant brain activations in response to images at 6 Hz up to 24 Hz. At 100% contrast only, face responses are visible at the 1.2 Hz component and its harmonics up to 16.8 Hz. No significant face response is observed at 0% contrast.

In addition, face images elicited strong responses at 1.2 Hz and harmonics at all pre-defined regions-of-interest (ROI 1 to ROI 5), which suggested that participants’ brains automatically detected their appearance every 5^th^ item. At 1.2 Hz, these responses were significant (z-score threshold = 3.09, p < .001, 1-tailed) at the medial occipital cortex (ROI 1: mean SNR = 2.58; electrodes’ z-scores at 1.2 Hz > 8.18) and progressively increased in magnitude as moving forwards through the right sensor space (ROI 2: mean SNR = 3.34; electrodes’ z-scores at 1.2 Hz > 12.51; ROI 4: mean SNR = 3.36; electrodes’ z-scores at 1.2 Hz > 8.76) and the left sensor space (ROI 3: mean SNR = 3.74; electrodes’ z-scores at 1.2 Hz > 12.25; ROI 5: mean SNR = 3.37; electrodes’ z-scores at 1.2 Hz > 7.30), without any statistical difference between the two sides of the scalp (ROI 2 *vs*. ROI 3: 2-tailed paired t-test: t(14) = −.793, p = .441; ROI 4 *vs*. ROI 5: 2-tailed paired t-test: t(14) = .110, p = .914). Images at 0% (i.e., dark grey squares flickering on a light grey background) did not elicit any significant face response, at none of the ROIs (z-score threshold = 3.09, p > .001).

These preliminary findings thus confirm that participants’ SSVEP measurements in response to faces and their visibility ratings, as assessed through two visibility scales adapted to SSVEP stimulation, can be associated without contaminating each other. As such, they provide new opportunities to explore people’s phenomenology when they see complex visual images such as faces.

### (3) Subliminal activation and propagation of the face signal along a posterior-to-anterior axis

When looking at the scalp topographies generated in response to faces across conditions, we observed two distinct phenomena. First, the signal propagated along a posterior-to-anterior axis, bilaterally, as face contrast increased (left: F(2,28) = 14.30, p < .001; right: F(2,28) = 12.82, p < .001; Figure 4A). Second, its modulation was more important over posterior than anterior brain regions, as suggested by the observation of larger modulation between ROI 2 and ROI 4 on the right side (t(14) = −3.594, p = .003) and ROI 3 and ROI 5 on the left side of the scalp (t(14) = −4.340, p < .001) than between ROI 1 and ROI 2 on the right (t(14) = −1.090, p = .294) and ROI 1 and ROI 3 on the left (t(14) = −1.255, p = .230) (Figure 4B).

**Figure 4.**
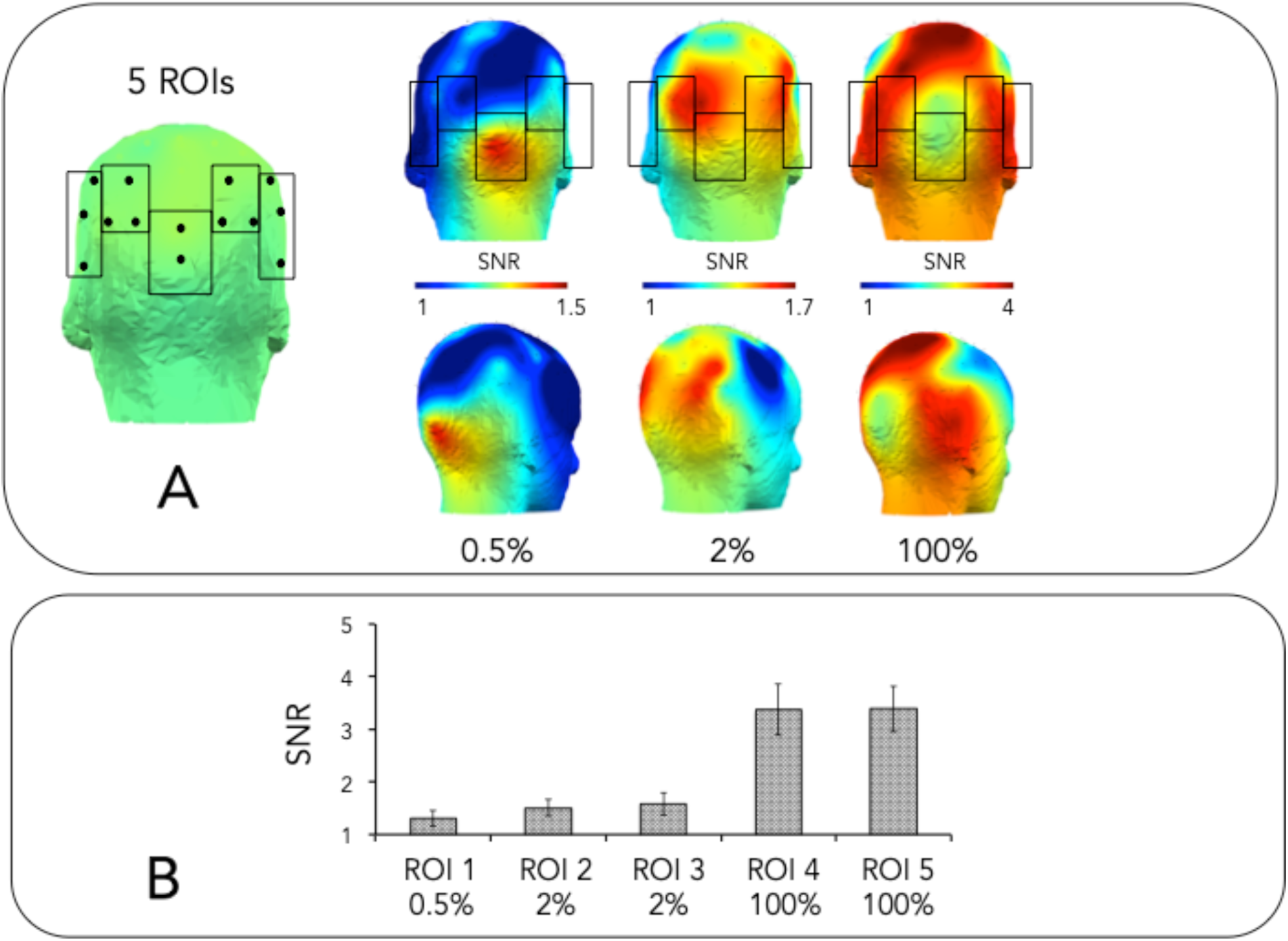
Propagation of the SSVEP signal along a posterior-to-anterior axis targeted through 5 regions-of-interest (ROI). The face signal extracted at 1.2 Hz was confined to a very posterior zone in response to subliminal images (ROI 1 capturing face response at 0.5% contrast - max SNR = 1.5) and propagated along a posterior-to-anterior axis, bilaterally, towards ROI 2 and ROI 3 first, which best captured the face response at 2% contrast (max SNR = 1.7) and further to ROI 4 and ROI 5, which best captured the face response at 100% contrast (max SNR = 4). At this latter contrast level, the activation of central electrodes was visible at 1.2 Hz but not observed at subsequent harmonics. B. The face signal also increased in amplitude along this axis. Bars and error bars represent group performance and standard errors, respectively.

When refining the story for each contrast level separately, we observed that, at 2% contrast, the face response extracted at 1.2 Hz peaked significantly (z-score threshold at 3.09, p < .001, 1-tailed) within ROI 2 at electrode PO8 (SNR at 1.2 Hz = 1.61 (SE = .19); z-score = 3.19) and within ROI 3 at electrodes O1 (SNR at 1.2 Hz = 1.67 (SE = .26); z-score = 3.52) and PO3 (SNR at 1.2 Hz = 1.58 (SE = .23); z-score = 3.15). It did also not leak down to ROI 4 or ROI 5 (Figure 4A) or to other harmonics, to the exception of a significant face response at 2.4 Hz at PO3 electrode. At lower contrast levels, namely at 1% and 1.5% contrast, the 1.2 Hz face response did not reach significance at any of the ROIs (see discussion). At last, the face response recorded at 0.5% contrast reached significance but, interestingly, only at ROI 1. Complementary analyses indicated that the only electrode responding significantly to the repetition of a face item every 5^th^ element with the same statistical constraints (z-score threshold at 3.09; p < .001, 1-tailed) as those imposed at 2% contrast was in fact Iz located at the level of the primary visual cortex (SNR = 1.52 (SE = .19); z-score = 3.83) (Figure 4A). Impressively, the magnitude of its response was not statistically different from the face response recorded at O1 at 2% contrast (t(14) = −.488, p = .633) and did also not spread to other harmonics.

### (4) Posterior brain activation predicts phenomenal experience

So far, we have observed that the amount of contrast embedded in the faces to which participants were exposed was highly predictive of the scalp topography at which the response was maximally observed. The current SSVEP paradigm can nevertheless only be considered as a no-report paradigm if some neural signatures such as the amplitude of the signal and/or the scalp topography predicts phenomenal experience.

To test this hypothesis, we focused on the contrast levels at which significant brain activation in response to faces could be delineated (0.5%, 2% and 100%) and explored, for each of these contrast levels separately, whether participants’ SNR recorded at their underlying ROIs (ROI 1 for 0.5% contrast - ROI 2 and ROI 3 for 2% contrast and ROI 4 and ROI 5 for 100% contrast) could predict their visibility ratings collected at SESSION 2. As qualitative and quantitative visibility ratings did not correlate to one another, at none of the contrast levels (one-tailed Pearson correlations: rs > .05), we considered them separately in the analyses. Qualitative ratings also reached ceiling values at 100% and were therefore not included in the analyses. As such, calculations involved, a series of correlation/regression analyses, all corrected for multiple comparisons.

Correlational analyses were only conclusive at 2% contrast. At this contrast level, participants’ averaged SNR in response to faces correlated moderately to their quantitative visibility ratings, indexed by the number of images they reported having seen, both at ROI 2 (1-tailed Pearson correlation: r = .421, p = .059) and ROI 3 (1-tailed Pearson correlation: r = .482, p = .035). None of these correlations survived the Bonferroni correction applied for multiple comparisons (i.e., 3 for the qualitative scale and 5 for the quantitative scale).

Given that we were interested in the directionality of the objective-subjective relationship, we decided to run complementary regression analyses on the signal recorded at 2% contrast and used the electrodes of ROI 2 and ROI 3 separately as predictors of participants’ qualitative and then quantitative visibility ratings. For these analyses, we favoured the backward method so that all predictors could be entered into the equation and then deleted one at the time to evaluate their individual contribution to the regression equation. Three and 5 regression analyses were computed on qualitative and quantitative ratings, respectively. Their observed alpha values were all corrected for multiple comparisons.

From the analyses performed, the model appeared to be predictive of participants’ quantitative, but not qualitative, ratings. This was especially true for quantitative ratings when O2 and PO8 electrodes were introduced in the computation. This reduced set of predictors indeed accounted for nearly as much of the variance (R^2^ = .429) as what was accounted for when all 3 electrodes composing ROI 2 were included in the analyses (PO4, PO8 and O2; R^2^ = .443), but this time significantly (F(2,14) = 4.508, p = .035). The size of the effect increased even more when P08 only was considered in the analyses (R^2^ = .344; F(1,14) = 6.811, p = .022; see Figure 5A for the corresponding correlational analysis). The same model but computed for the left part of the scalp, and therefore on the electrodes of ROI 3, revealed significant results but only when electrode O1 was kept into the analyses (F(1,14) = 4.902, p = .045). The proportion of variance this model explained (R^2^ = .274) was however not as high as the one of the full model (R^2^ = .355) or the one observed for the right part of the scalp (R^2^ = .429).

**Figure 5.**
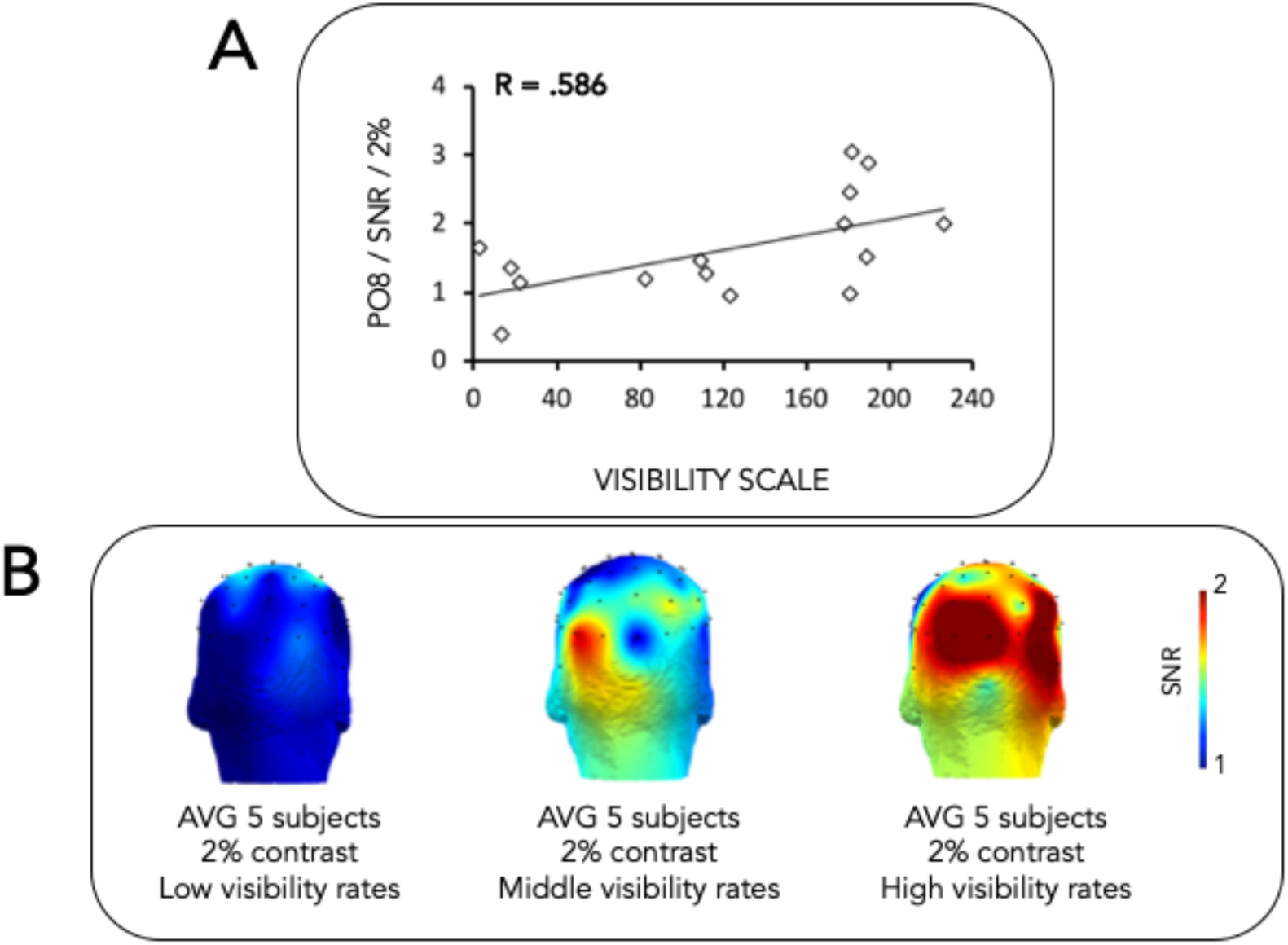
Amplitude of participants’ SSVEP response to faces and their corresponding scalp topography at 2% contrast being indicative of the fact that the SSVEP tool could potentially be used in the future as a no-report paradigm to capture phenomenal experience. A. Participants’ face categorization responses (1.2 Hz; N=15) collected at PO8 positively correlate with their visibility judgement of how many images they estimate to have categorized. B. Their scalp topographies are also indicative of their phenomenal experience.

Anecdotally, it was also interesting to observe that the brain regions participants recruit at 2% contrast was indicative of their phenomenology (Figure 5B). Participants who rated more images as visible at this contrast level were also those whose scalp topographies were most similar to the topographies observed at 100% contrast. As such, we would suggest that this contrast level could be taken as a reference point in the future when exploring whether the SSVEP outputs could serve as objective measures of subjective experience. At this fixed physical input, the objective SSVEP markers (amplitude of the SSVEP response to faces and corresponding scalp topography) seems indeed to predict participants’ subjective experience the best. Future studies could involve testing whether the same observation can be applied to other contrast levels.

## DISCUSSION

The goals of this study were threefold. First wanted to explore the extent to which complex visual images, such as faces, can be processed in the absence of consciousness and, second, to evaluate how their underlying signal propagates along a posterior-to-anterior axis as image contrast increases. We also wanted to assess whether a paradigm that does not require participants to give any overt (verbal or motor) response can be used to access their phenomenology. To address these questions, we tested a group of participants in two consecutive testing sessions requiring them to view the same set of complex visual images (faces, non-faces) varying systematically in terms of their overall contrast. The main goal of SESSION 1 was to establish, through a staircase procedure as well as through subjective reports, their visibility thresholds. Given the results obtained at the group level, images at 0.5% contrast were taken as subliminal. The main goal of SESSION 2 was to explore the extent to which the brain is sensitive to the periodic presentation of face images when their visibility varies. To do so, we compared participants’ brain activations in response to faces embedded in sequences of non-face stimuli. Stimulus contrast in these sequences also progressively increased over the course of the experiment. We predicted we would find a neural marker of face categorization at contrast levels for which participants reported having seen the stimuli (2% contrast and above), which would indicate the ability of their brains to categorize faces and differentiate them from non-face objects at supraliminal levels, even if their visibility had been degraded. Given the nature of the stimuli, namely faces, and of the SSVEP paradigm used here, we also expected to find a response in the brain even when reaching the threshold for which participants report having not seen the stimuli. Finally, two visibility scales (qualitative, quantitative) were provided to participants with the goal of assessing the visibility of the images embedded in SSVEP sequences to which they had just been exposed.

We think the results of this study are notable for three main reasons.

The first reason is that they document the exceptional capabilities of the human brain even in experimental conditions for which participants report the images as being unseen. Because of the power and sensitivity provided by the SSVEP technique, we were able to show that, over 6-7 minutes of stimulation only, the brain is not only capable of processing but also of creating a visual category from very different faces, even at a stage that had been defined in an earlier session as subliminal. This is spectacular given the very low visibility and high complexity of the stimuli that were involved in the task. Similar observations have been made in the past, but with stimuli of much lower complexity (e.g., gratings [19]; letters [34]) and in other contexts (e.g., working memory [35-36]). Interestingly, the face response extracted at the subliminal stage (0.5% contrast) was peculiar in the sense that it showed both harmonic and spatial restriction. Harmonic restriction relates to the fact that the face signal did not propagate significantly beyond 1.2 Hz, which corresponds to the frequency associated with the brain’s automatic response to the introduction of faces every 5^th^ item in the sequence. Such an observation is consistent with the finding that stimuli of low visual complexity typically engage fewer harmonics than those of high visual complexity [17,37]. Consistently with previous findings that unconscious information remains locally active in the brain, while it is made globally available to multiple brain systems once information becomes conscious [22, see also 41-43], spatial restriction was also obvious here in that the significant signal captured in response to subliminal faces remained confined to a single electrode located at the medial occipital lobe, which is typically activated by low-level visual information such as the orientation of lines embedded in the stimulus. This finding raises however the question of how such a brain activation can nevertheless be observed in response to complex visual stimuli such as faces. According to us, there are two non-mutually exclusive explanations. The first one is the involvement of invisible re-entrant feedback information originating from higher-level regions onto the medial occipital lobe [44-45]. The second one is that we used for the current study a subset of apparently heterogeneous faces but which, in fact, could have also engage repetitive structures (e.g., roundish shape) the striate cortex could have detected. To disentangle these proposals, one could test another group of participants on the same set of subliminal images but presented this time at inverted orientation. The low-level visual explanation would only be retained if medial occipital activations are also observed in response to inverted subliminal faces.

It is also worth noting that we did not observe any significant face activation at 1% and 1.5% contrasts. We attribute this observation to the fact that, in these conditions, participants started to allocate their attention to stimuli other than faces within the sequences, which inevitably impacted the size of the peaks negatively [37]. In line with this assumption is the observation that their visibility ratings also increased in these two conditions compared to the 0.5% contrast condition, even though not significantly. Future studies are needed to confirm this hypothesis.

The second reason for these results being notable is that beyond this point, namely at 2% contrast and above, we found that the face signal increased in amplitude as the image contrast increased and moved away from ROI 1, first to ROI 2 and ROI 3 and then to ROI 4 and ROI 5, namely those regions-of-interest underlying maximal activation in response to faces at 2% and 100% contrast. The current finding however raises the question of why we did not observe any (pre-)frontal activation in response to faces, even at 100% contrast. To us, this result should not be taken as evidence that this brain region had not been recruited at all during the task. Its activity could have indeed been captured at other frequencies than those associated with the repetitive introduction of faces within sequences or could even have been spread indistinctly over the frequency spectra if this part of the brain had been recruited during the whole task. Future studies could involve testing whether subjects show (pre-)frontal activation in response to faces when they have to explicitly report on their periodicity. This would help disentangling what is related to report, on the one side, and to phenomenology, on the other side.

The third reason for which these results are interesting is that they provide some evidence that SSVEP measurements can be used to predict participants’ phenomenology of complex visual images such as faces. This is not to undermine when thinking about vulnerable populations such as infants or some patients who cannot report on their phenomenology. In this experiment, we made the observation that the magnitude of participants’ brain responses recorded at electrodes typically engaged during face perception [15-16] were good predictors of phenomenal experience. We cannot however exclude the possibility that this variation in subjective vision is in fact due to individuals differing in their visual sensitivity since calibration has been performed at the group level. At a fixed contrast level, the present findings also emphasize the benefit of relying on a quantitative rather than a qualitative visibility scaling measure, which better fits the neuroimaging tool of the type of SSVEP used here. We would however recommend to explore the precise nature of the quantitative scale in depth in the future since its outputs do not correlate to those of the qualitative scale [46].

Overall, this study is part of a larger attempt to shed light on the neural correlates associated to the ability to categorize faces when they vary in terms of contrast and the phenomenology subjects can make of this experience by means of a powerful SSVEP paradigm derived from electroencephalography (EEG). We found that subliminal faces were significantly categorized at the level of the medial occipital lobe and that the magnitude of this categorization response increased as face contrast increased. Increasing contrast also drove different electrodes being significantly activated along a posterior-to-anterior axis on the scalp reminding of the visual tracks present in the brain. We finally observed that, at some specific scalp locations, topography and/or signal amplitude predicted participants’ phenomenology, which seems to highlight the potential of identifying the neural correlates associated with the experience of complex visual stimuli with greater precision in the future.

## ACKNOWLEDGMENTS

This research has been funded by an ERC advanced grant (RADICAL) attributed to AC in 2013. AC is Research Director is Research Associate of the Fonds de la Recherche Scientifique – FNRS, Belgium. We would like to thank Genevieve Quek for earlier comments on the manuscript and Fabienne Chetail for writing the script used for SSVEP analyses (https://osf.io/53hdb/; doi: 10.17605/OSF.IO/53HDB).

Data available on OSF (https://osf.io/hg6sm/).

## COMPETING INTERESTS

The authors declare that no competing interest.

## AUTHOR CONTRIBUTIONS

A.d.H. designed the experiment, carried out the analyses and wrote the paper. LS tested the participants. AB, LV and AC wrote the paper.

## Notes

### Competing Interest Statement

The authors have declared no competing interest.

